# Transcriptomic analysis of primate placentas and novel rhesus trophoblast cell lines informs investigations of human placentation

**DOI:** 10.1101/2020.08.21.262030

**Authors:** Jimi L. Rosenkrantz, Jessica E. Gaffney, Victoria HJ. Roberts, Lucia Carbone, Shawn L. Chavez

**Author notes:** To whom correspondence should be addressed: Shawn L. Chavez, Ph.D., 505 NW 185^th^ Avenue, Beaverton, OR 97006, Lucia Carbone, Ph.D., 3303 SW Bond Avenue, Portland, OR 97239.

## Abstract

Proper placentation, including trophoblast differentiation and function, is essential for the health and well-being of both the mother and baby throughout pregnancy. Placental abnormalities that occur during the early stages of development are thought to contribute to pre-eclampsia and other placenta-related pregnancy complications. However, relatively little is known about these stages in humans due to obvious ethical and technical limitations. Rhesus macaques are considered an ideal surrogate for studying human placentation, but the unclear translatability of known human placental markers and lack of accessible rhesus trophoblast cell lines can impede the use of this animal model. Here, we performed a cross-species transcriptomic comparison of human and rhesus placenta and determined that while the majority of known placental markers were similarly expressed, 952 differentially expressed genes (DEGs) were identified between the two species. Pathway enrichment analysis of the 447 human-upregulated DEGs, including *ADAM12*, *ERVW-1*, *KISS1*, *LGALS13*, *PAPPA2*, *PGF*, and *SIGLEC6*, revealed over-representation of functional terms associated with pre-eclampsia and other pregnancy disorders. Additionally, to enable *in vitro* functional studies of early placentation, we generated and thoroughly characterized two highly-pure first-trimester telomerase (TERT) immortalized rhesus trophoblast cell lines (iRP-D26 and iRP-D28A) that retained crucial features of isolated primary trophoblasts. Overall, our findings help elucidate the molecular translatability between human and rhesus placenta and reveal notable expression differences in human placental markers and genes associated with pregnancy complications that should be considered when using the rhesus animal model to study normal and pathological human placentation.

## Introduction

The placenta is the physical link between the mother and fetus as well as a critical site for nutrient and waste exchange during pregnancy. Trophoblasts are the major functional cell type of the placenta, and can be divided into three subtypes: (1) proliferative cytotrophoblasts (CTBs), which continually fuse to form (2) multinucleated syncytiotrophoblasts (STBs), and (3) invasive extravillous trophoblasts (EVTs) that remodel the uterine spiral arteries to gain an adequate blood supply. Proper trophoblast differentiation and function is essential for normal placentation and fetal development throughout gestation. In humans, abnormal placentation is the primary defect associated with major pregnancy complications, such as pre-eclampsia, fetal growth restriction, recurrent miscarriage, and still-birth (Brosens et al. 2011). Investigation of early placentation is particularly important for combating these diseases since many of the associated defects are thought to arise early to mid-gestation. However, the ethical and technical limitations of studying early human development have confined many human placentation investigations to late gestational stages closer to term. Thus, early human placental development, including the origin and cause(s) of the placental abnormalities underlying major pregnancy complications, are poorly understood.

To overcome the limitations of studying early human placentation, numerous human first-trimester trophoblast cell lines have been isolated and immortalized using various methods for *in vitro* investigations (Pattillo and Gey 1968; Graham et al. 1993; Straszewski-Chavez et al. 2009). Unlike primary trophoblasts, immortalized trophoblast cell lines are readily available, can be grown in culture indefinitely, and are relatively easy to transfect for functional studies. However, recent studies suggest that these cell lines are not necessarily a pure population of trophoblasts and/or have acquired karyotypic and phenotypic aberrations with continued passaging (Apps et al. 2009; Abou-Kheir et al. 2017; Reiter et al. 2017). Moreover, all of the established cell lines are EVT in origin (Abbas et al. 2020), which suggests that additional sources of CTBs are required to study trophoblast differentiation and syncytialization. The human choriocarcinoma cell lines, BeWo, JEG-3 and JAR, have also been used for this purpose (Kohler et al. 1971), but these cells are highly malignant and exhibit considerably different transcriptomic profiles than primary trophoblasts (Apps et al. 2009), questioning whether findings using these lines are truly representative of normal CTBs or EVTs.

While some of the limitations of performing human *in vitro* and *in vivo* placental studies could be surmounted using animal models, most mammalian species poorly recapitulate human placentation. This is due, in large part, to inherent genetic differences and variations in the placental structure, steroidogenic synthesis, and the intracellular signaling pathways mediating lineage-specific trophoblast differentiation amongst mammals (Malassine et al. 2003; Soncin et al. 2015). However, non-human primate animal models, particularly rhesus macaques (*Macaque mulatta*), are genetically similar to humans and share many key features of human placentation. Besides being comparable in placental morphogenesis, the overall structure and nature of both the STB interface layer and intervillous space, as well as endocrine functions and extracellular matrix changes are similar between rhesus and human placenta (Ramsey and Harris 1966; Ramsey et al. 1976; Ellinwood et al. 1989; King and Blankenship 1994). Further, there is a strong resemblance between human and rhesus placental endovascular EVT invasion and spiral artery transformation (Blankenship and Enders 2003), processes known to play a central role in the pathogenesis of pre-eclampsia and fetal growth restriction in humans (Burton et al. 2009). Unlike placental studies relying on human samples, access to rhesus early gestational placental samples and *in vivo* functional investigations are possible under approved animal studies. However, rhesus first-trimester trophoblast cell lines are still not readily available, which limits rhesus placental studies and requires the laborious isolation and use of primary term rhesus trophoblasts for *in vitro* investigations.

Although the human and rhesus placenta appear to be morphologically and functionally similar, previous studies have revealed some notable differences in the expression level and/or protein-coding potential of the well-known human placental markers, including CGA, HLA-G, ERVW-1, and SIGLEC6 (Maston and Ruvolo 2002; Boyson et al. 1996; Esnault et al. 2013; Brinkman-Van der Linden et al. 2007). Differences in placental invasion have also been noted, as the extent and depth of interstitial EVT invasion is greater in human compared to rhesus placentation (Ramsey et al. 1976). Further, while a few cases of pre-eclampsia have been documented in rhesus and other non-human primates, this disease predominantly occurs in humans (Krugner-Higby et al. 2009; Palmer et al. 1979; Stout and Lemmon 1969; Carter 2007; Abbas et al. 2020). Thus, the identification of the molecular differences between human and rhesus placenta will not only help elucidate the translatability between primate placental studies, but it may also provide valuable insight into the molecular origin of human-specific placental features and pregnancy-related diseases. The lack of comprehensive transcriptome comparison of human and rhesus placenta currently inhibits our understanding of these differences at a molecular level. Here, we performed a cross-species transcriptomic comparison of human and rhesus placenta to identify DEGs and determined that even though the majority of placental markers are similarly expressed across the two species, there are gene expression differences that likely underlie the distinct features of human placentation. Additionally, we generated and thoroughly characterized two highly-pure TERT-immortalized rhesus trophoblast cell lines for *in vitro* functional studies that retained features of primary rhesus trophoblasts. Overall, this work provides a comprehensive list of genes differentially expressed between human and rhesus placenta that enhances the translatability of primate placental investigations, helps delineate the molecular differences underlying human susceptibility to pre-eclampsia and other pregnancy-related diseases, and provides previously unavailable first-trimester rhesus trophoblast cell lines for further functional investigation and understanding of early primate placentation.

## Results

### Identification of genes differentially expressed between human and rhesus placenta

Despite the structural and functional similarities between human and rhesus placentas, differences in the level and route of invasion, as well as the increased propensity to develop pregnancy-related diseases in humans, suggests that molecular differences exist. To characterize such differences, we compared gene expression levels between human and rhesus macaque (*Macaque mulatta*) placentas using a combination of existing and newly-generated RNA-seq data from bulk term placental tissues (Dunn-Fletcher et al. 2018) (Supplemental Table S1). Given the challenges of performing cross-species differential expression (DE) analyses, DEGs were identified by combining two approaches (Supplemental Fig. S1A). First, RNA-seq data from both species were mapped to the human reference genome (GRCh38) (Fig. 1A) and DEGs were identified based on the human gene annotations (DE-GRCh38). Second, RNA-seq data from both species were mapped to the rhesus reference genome (Mmul10) and DEGs were identified based on the rhesus gene annotations (DE-Mmul10) (Fig. 1B). A gene was considered differentially expressed only if it was identified as significantly (padj<0.05) upregulated or downregulated (|L2FC|>2) by both analyses (Fig. 1C). In order to avoid potential batch effects, we repeated the DE analysis using three independent human placenta RNA-seq datasets (Green et al. 2016; Pavlicev et al. 2017) (Supplemental Fig. S1A-C). The final set of DEGs consisted only of genes determined to be significantly upregulated or downregulated by all three DE analyses and resulted in a total of 952 DEGs, including 447 human-upregulated and 505 rhesus-upregulated genes (Fig. 1D, Supplemental Table S2). Eight of the DEGs were validated using a quantitative reverse transcription PCR (qRT-PCR) approach (Supplemental Fig. S2A,B). The 25 most significant human and rhesus upregulated DEGs are shown in Figure 1E and highlights the presence of several well-known placental markers upregulated in human (*ADAM12*, *SERPINB2*, *BPGM*, *CYP19A1*, *SVEP1*, *GPC3*, *PGF*, *FBN2*, and *PAPPA2)*, several of which are associated with highly-invasive human EVTs (Biadasiewicz et al. 2014; Aghababaei et al. 2014; Wang et al. 2016). Pathway enrichment analysis revealed that functional terms related to “extracellular region/space” were uniquely over-represented in the rhesus-upregulated gene set, while functional terms associated with pregnancy disorders including, “pre-eclampsia” (padj=6.73E-04), “HELLP syndrome” (padj=1.48E-01), “gestational trophoblastic tumor” (padj=1.52E-01), and “eclampsia” (padj=4.43E-01) were uniquely over-represented in the human-upregulated gene set (Supplemental Fig. S3, Supplemental Table S3,S4). These results provide a comprehensive list of DEGs between human and rhesus placenta and indicate that certain well-known human placental/trophoblast cell markers may not have equivalent expression patterns in rhesus placenta. They also suggest that human-upregulated genes are associated with human-specific placental features, which may contribute to the heightened occurrence of pre-eclampsia and other pregnancy-related diseases in humans.

**Figure 1.**
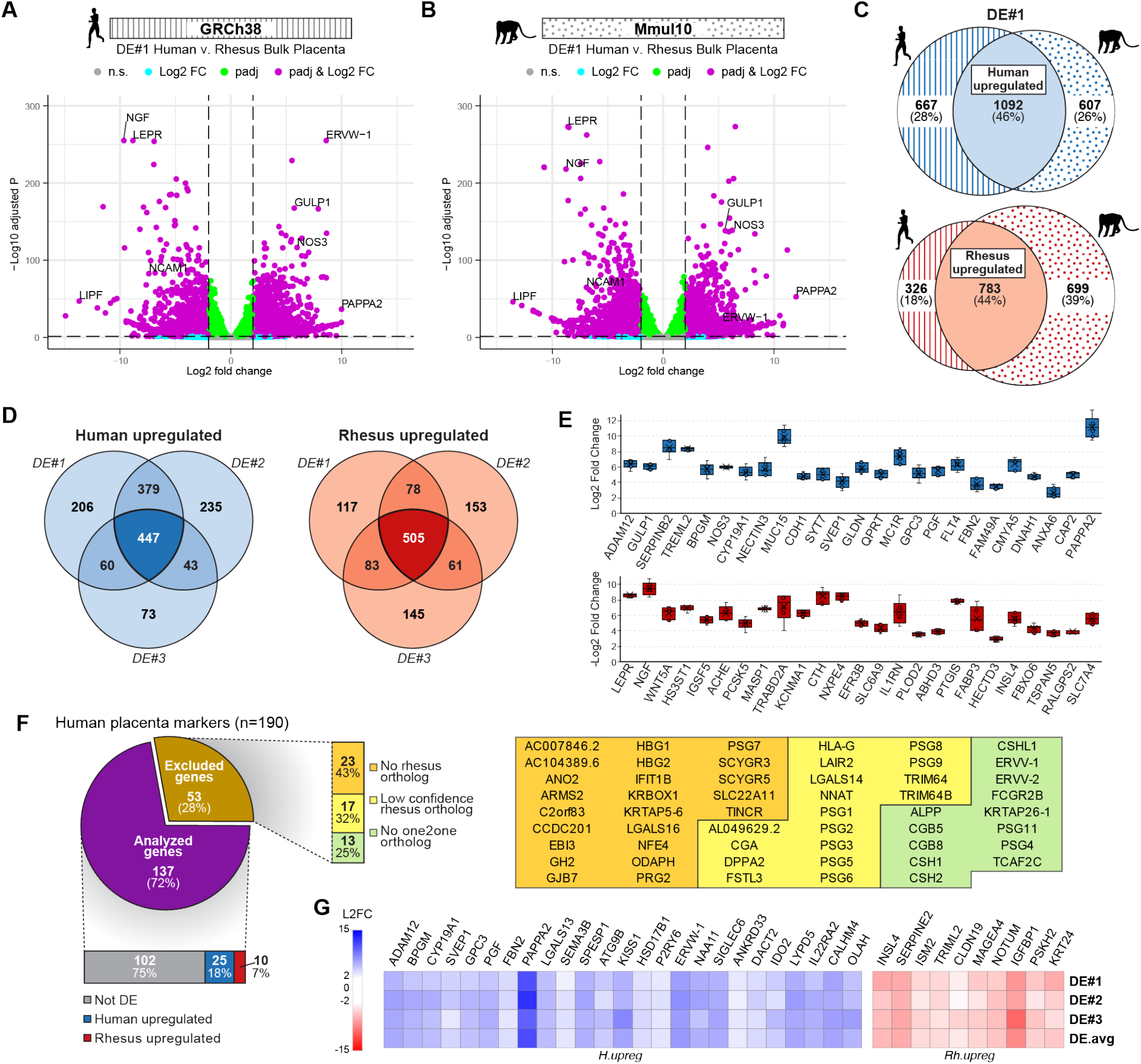
Cross-species transcriptional comparison of human and rhesus bulk placental tissue. Volcano plots showing gene expression fold differences between human and rhesus term placental tissue from DE#1, using data mapped to (A) the human genome and (B) the rhesus genome. Dashed lines denote DE significance (padj<0.05) and fold change (|L2FC|>2) thresholds; genes passing padj threshold (green), passing L2FC threshold (cyan), passing both (magenta), or none (grey). (C) Venn-diagram depicting intersection of DEGs identified using data mapped to human genome (stripes) and rhesus genome (spotted) to identify intermediate human-upregulated (light blue) and rhesus-upregulated (light red) genes sets. (D) Venn-diagram depicting the intersection of the results from the three DE analyses to identify final set of 447 human-upregulated genes (blue), and 505 rhesus-upregulated genes (red). (E) Top 25 most significant differentially expressed (ranked by mean padj) human-upregulated (blue) and rhesus-upregulated genes (red); box plots depict average Log2 fold change of each gene from the three DE analyses. (F) Examination of human placental marker genes DE results. (Left) Proportion of placental marker genes analyzed (purple) or excluded from (brown) DE analysis. Analyzed genes are further classified as either not differentially expressed (not DE) (grey), upregulated in human (blue), or upregulated in rhesus (red). (Right) Classification of excluded human placental marker genes with no rhesus ortholog (orange), with low confidence rhesus ortholog (yellow), or lacking a one2one rhesus ortholog (green). (Bottom) Heatmap of differentially expressed placental markers showing expression level changes in human compared to rhesus placenta; human-upregulated genes (L2FC>2) are shown in blue and rhesus-upregulated genes (L2FC<2) are shown in red.

### Comparison of placental marker expression between human and rhesus placenta

To elucidate the translatability of human placental marker gene expression to rhesus placenta, a set of previously defined human placental markers (Uhlén et al. 2015) was examined. Out of the 190 placental markers, 72% (N=137/190) were included in the DE analysis, while 28% (N=53/190) were excluded due to nonexistent (43%), low confidence (32%), or a lack of “one2one” (25%) rhesus ortholog (Fig. 1F). Of the 137 placental markers included in the analysis, the majority (~75.5%; N=102/137) showed similar expression levels and were not differentially expressed between human and rhesus placenta. The remaining genes (~25.5%; N=35/137) were identified as differentially expressed, with ~18% (N=25/137) being upregulated in human and ~7% (N=10/137) upregulated in rhesus (Fig. 1F, Supplemental Table S5). Notably, placental markers associated with invasive EVTs (*ADAM12, LGALS13, PAPPA2*, *PGF*) (Hutter et al. 2016; Knuth et al. 2015; Athanassiades and Lala 1998), and pregnancy complications such as preterm birth and pre-eclampsia (*ADAM12, HSD17B1, KISS1, LGALS13, PAPPA2, SIGLEC6, ERVW-1*) (Enquobahrie et al. 2012; Ishibashi et al. 2012; Kapustin et al. 2020; Sekizawa et al. 2009; Nishizawa et al. 2008; Kaartokallio et al. 2015; Knerr et al. 2002), were found to be upregulated in human compared to rhesus placental tissue (Fig. 1G). Additionally, the subset of placental marker genes lacking one-to-one rhesus orthologs included the EVT marker, *HLA-G* (Boyson et al. 1996), genes encoding different human chorionic gonadotropin (hCG) subunits (*CGA, CGB5*, and *CGB8*), chorionic somatomammotropins hormones (*CSH1, CSH2, CSHL1*), and pregnancy-specific glycoproteins (*PSG1–PSG9*, *PSG11*). Nonetheless, the vast majority (75.5%) of human placental marker genes were similarly expressed between human and rhesus, indicating that rhesus placenta can serve as a suitable surrogate for studying most features of human placentation. However, the gene expression differences identified here should be considered when studying certain aspects of placentation in rhesus, particularly those that are associated with EVT invasion and pre-eclampsia or other placenta-related diseases.

### Establishment of TERT-immortalized rhesus placental and skin fibroblast cell lines

Because the placenta is a heterogeneous organ comprised of many cell types in addition to trophoblasts, such as immune, stromal and vascular cells (Liu et al. 2018), we next sought to isolate primary trophoblasts from bulk rhesus placentas for immortalization and characterization. Trophoblast cells were isolated from both first and third-trimester rhesus placentas since there have only been reports of successful rhesus term trophoblast cell derivation (Golos et al. 2006). Using a strategy described in Figure 2A, we isolated primary trophoblast cells from bulk rhesus placental tissues collected at gestational day 26 (~6 weeks human pregnancy), day 28 (~7 weeks human pregnancy), day 50 (~12 weeks human pregnancy), day 141 (~34 weeks human pregnancy), and day 149 (~35 weeks human pregnancy). After depletion of contaminating immune cells using immunopurification, the cells were cultured for 24 hours before transduction with lentivirus containing *TERT* and puromycin resistance (*PAC*) genes for antibiotic selection. A total of six immortalized rhesus placental (iRP) cell lines were generated, including four from first-trimester (iRP-D26, iRP-D28A, iRP-D28B, iRP-D50) and two from third-trimester (iRP-D140, iRP-D141) rhesus placentas. Male and female rhesus primary skin fibroblasts were also used to establish two TERT-immortalized rhesus fibroblast (iRFb) cell lines, iRFb-XY and iRFb-XX, as controls. Cultures of iRP-D26 and iRP-D28A contained purely polygonal epithelial-like cells, while the other cell lines appeared heterogeneous with a mix of large flattened and elongated fibroblast-like cells (Fig. 2B). Expression of the lentiviral-transduced genes, *TERT* and *PAC*, was confirmed in each of the cell lines via RT-PCR. (Fig. 2C).

**Figure 2.**
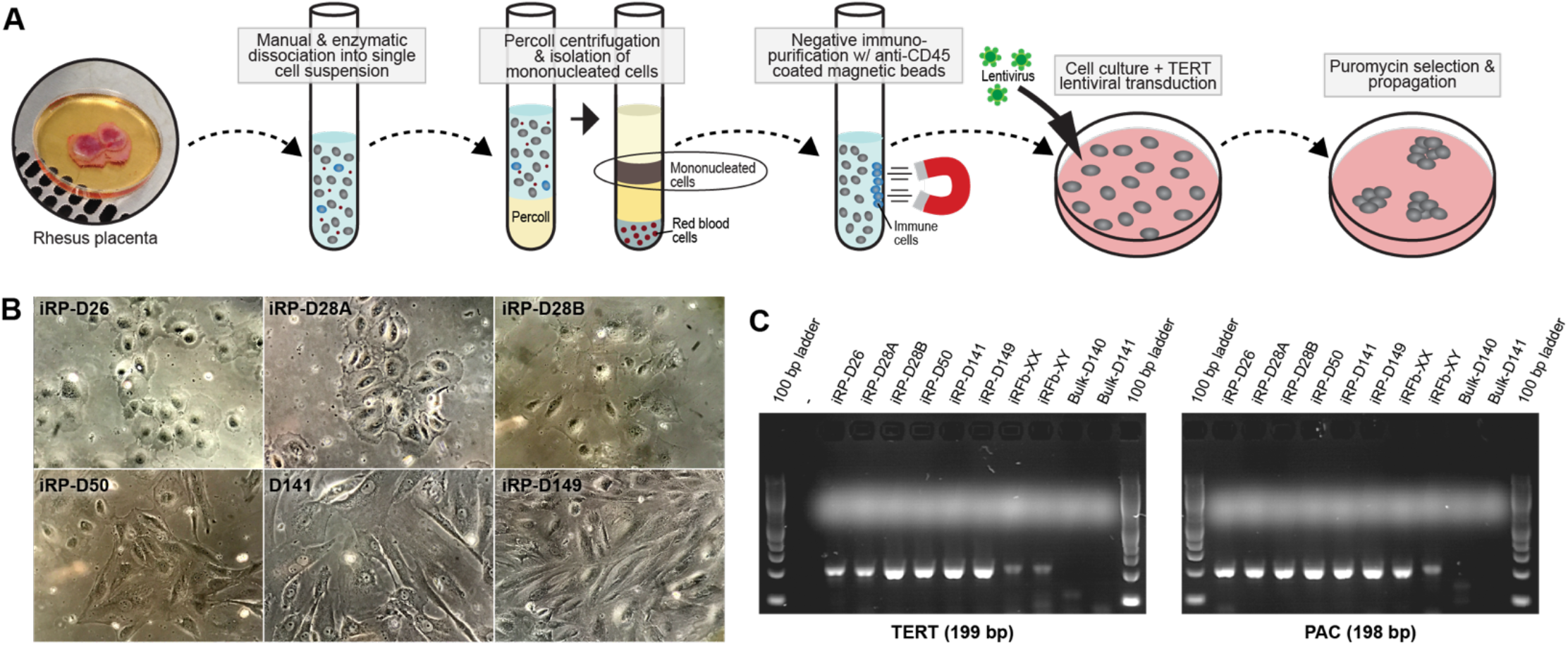
Establishment of TERT immortalized rhesus placental and skin fibroblast cell lines. (A) Schematic of primary trophoblast cell isolation from rhesus placental tissue and TERT immortalization. (B) Phase contrast microphotographs of immortalized placental cell lines. (C) Confirmation of TERT immortalization via RT-PCR detection of *TERT* and *PAC* gene expression.

### Assessment of genomic integrity in immortalized placental and fibroblast cell lines

Cell lines are known to develop genetic abnormalities during immortalization and/or prolonged cell culture, such as aneuploidies, copy number variations (CNVs), and chromosomal fusions. This is particularly true for cell lines derived by Simian Virus 40 (SV40) or a similar transformation approach as has been shown for first-trimester human trophoblast cell lines (Aboagye-Mathiesen et al. 1996; King et al. 2000), but it can also occur in TERT-immortalized human trophoblasts over time (Reiter et al. 2017). In addition, primary trophoblasts normally undergo cell fusion (syncytialization), which can complicate nuclear assessment. Here, CNVs and whole chromosome counts were determined using low-input DNA-seq and metaphase spreads, respectively. For each cell line, approximately ten cells were manually transferred into a single tube and prepared for DNA-seq as previously described (Daughtry et al. 2019). The normal number of diploid chromosomes (N=42) was detected in iRP-D26, iRP-D28A and iRP-D141, although sub-chromosomal losses of Chr1 and Chr7 were observed in iRP-D26, sub-chromosomal gains of Chr11 and Chr16 in iRP-D28A, and a small sub-chromosomal loss of Chr1 in iRP-D141 (Fig. 3A). In contrast, numerous whole and sub-chromosomal CNVs were identified in the other immortalized placental cell lines, iRP-D28B, iRP-D50, and iRP-D149 (Fig. 3A). As expected, CNV analysis of the female rhesus fibroblasts (iRFb-XX) showed normal diploid copy numbers for all 21 rhesus chromosomes, while the male fibroblasts (iRFb-XY) exhibited the expected ChrX “loss” and detection of ChrY (Fig. 3A, B). Comparison to the iRFb-XY control revealed a single copy of ChrX without the detection of ChrY in iRP-D26, highlighting the loss of a whole sex chromosome (Fig. 3A, B). Metaphase spreads confirmed the loss of one to two whole chromosomes in iRP-D26 cells, supporting the DNA-seq results, and suggesting the existence of chromosome fusion in cells with only 40 chromosomes (Fig. 3C, D). Examination of metaphase spreads from iRP-D28B, iRP-D50, and iRP-D149 further demonstrated a heterogenous mix of predominantly polyploid cells, containing between three (triploid) and four (tetraploid) sets of chromosomes (Fig. 3C, D).

**Figure 3.**
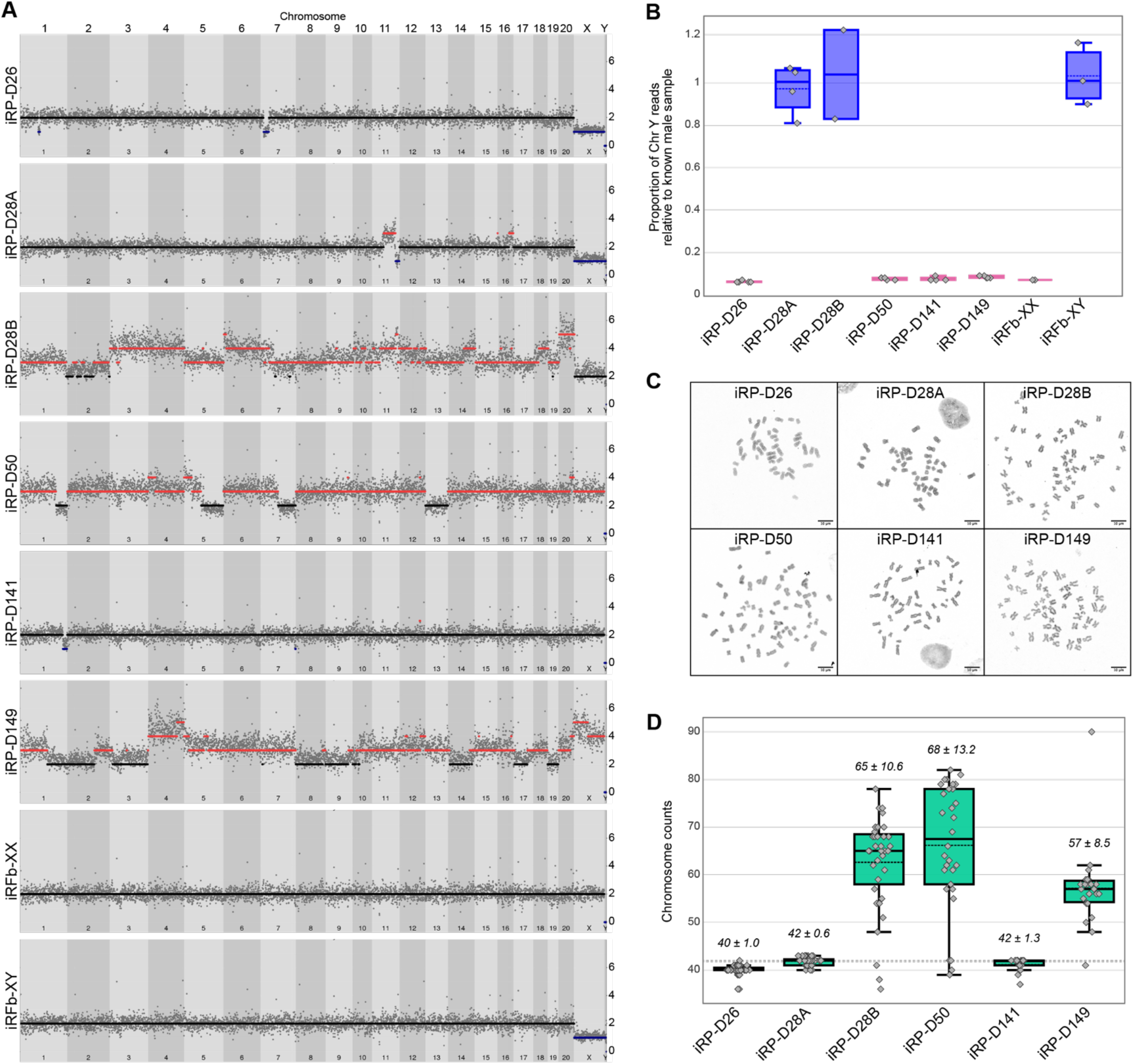
Assessment of genomic integrity in immortalized placental cell lines. (A) Manhattan plot of whole genome CNVs organized by chromosome; copy number gains (red), copy number losses (blue) show the presence of CNV in most of the placental cell lines but not in the fibroblast used for comparison. (B) Box plots depicting relative detection of ChrY, normalized to known male sample, iRFb-XY; cell lines were identified as male if mean > 0.5 (blue), or as female if mean < 0.5 (pink). (C) Representative metaphase spreads for each of the iRP cell lines generated; 10uM scale bar. (D) Box plots of metaphase spread chromosome count results; median ± standard deviation values included above each plot.

### iRP-D26 and iRP-D28A represent two highly pure immortalized rhesus trophoblast cell lines

While our trophoblast isolation procedure was optimized to reduce contamination of non-trophoblast cell types, low levels of contaminating stromal and immune cells may persist following immortalization, selection, and propagation. In order to identify the placental cell lines containing a pure population of trophoblast cells, we analyzed each line for expression of highly conserved trophoblast and non-trophoblast cell markers (Lee et al. 2016), including, KRT7, a pan-trophoblast cell marker; CDH1, a mononuclear trophoblast cell marker; VIM, non-trophoblast stromal marker; and PTPRC (CD45), a pan-leukocyte marker. Antibodies for these markers were first validated in rhesus placental tissues using immunohistochemical (IHC) staining and the observed expression patterns were consistent with known patterns in the human placenta (Fig. 4A). Immunofluorescent (IF) staining using the same antibodies showed robust staining of KRT7 and CDH1 trophoblast markers, and the absence of VIM and CD45 staining in iRP-D26 and iRP-D28A, indicating the enrichment of trophoblasts and the absence of mesenchymal and immune cells within these cell lines, respectively. In contrast, the remaining immortalized placental cell lines (iRP-D28B, iRP-D50, iRP-D141, and iRP-D149) were largely contaminated with VIM positive mesenchymal cells (Fig. 4B). These findings were consistent with qRT-PCR expression analysis, which showed significant enrichment of KRT7 and CDH1 expression in iRP-D26 and iRP-D28A compared to the other cell lines or bulk rhesus placental tissue (Fig. 4C). Additionally, VIM and PTPRC (CD45) expression was not detected by qRT-PCR in iRP-D26 and iRP-D28A, but were highly expressed in all other cell lines analyzed (Fig. 4C). Collectively, these results highlight the enrichment of mononuclear trophoblast and the absence of non-trophoblast cells in the iRP-D26 and iRP-D28A cell lines.

**Figure 4.**
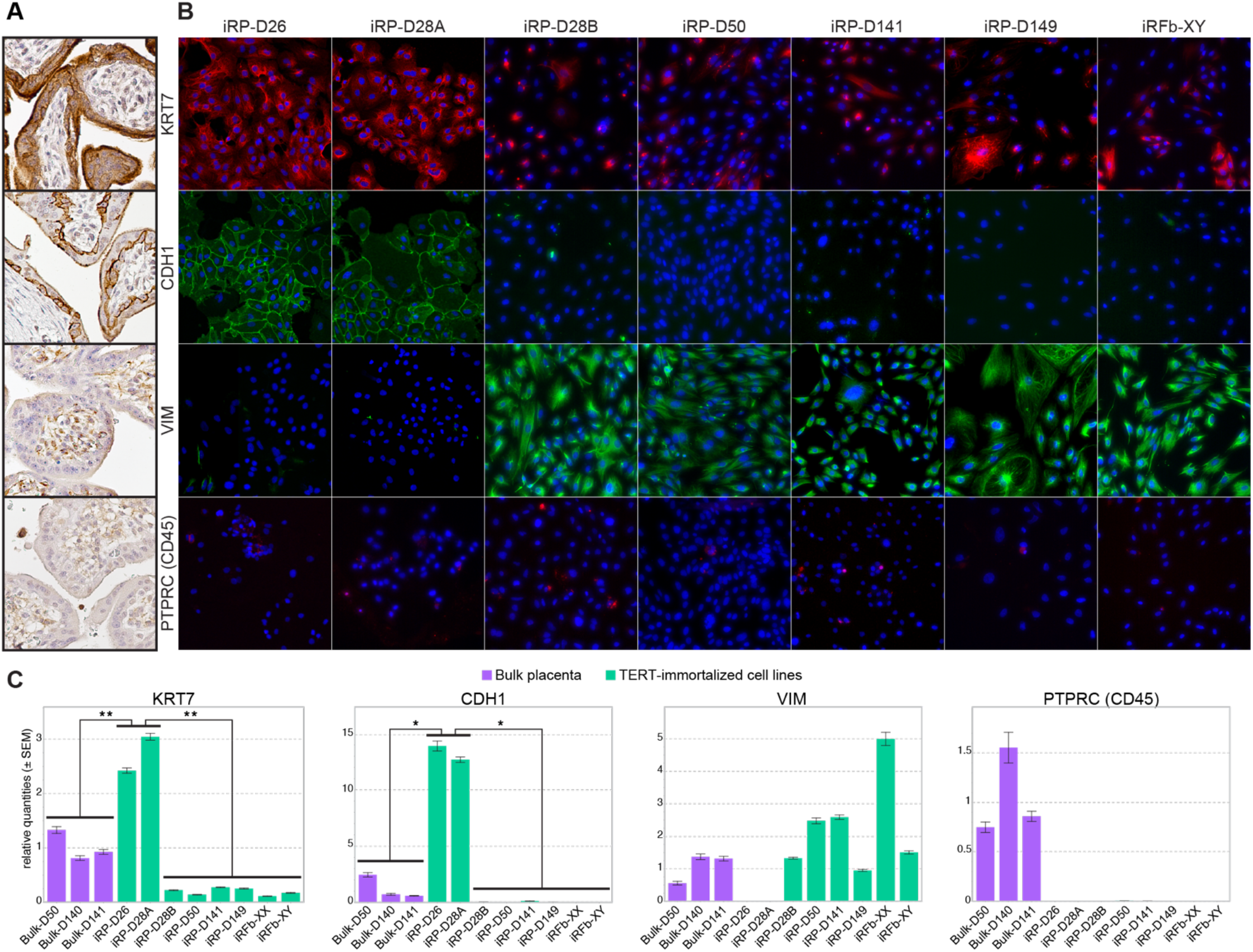
iRP-D26 and iRP-D28A represent two highly pure rhesus immortalized trophoblast cell lines. (A) IHC staining of gestational day 50 rhesus placental tissue for mononuclear trophoblast (KRT7 and CDH1), stromal (VIM), and immune cell (PTPRC) markers (DAB, brown); hematoxylin nuclear counter stain (blue). (B) IF staining of immortalized cell lines for KRT7 (red), CDH1 (green), VIM (green), and PTPRC (red); DAPI nuclear counterstain (blue); results show that iRP-D26 and iRP-D28A cells express known mononuclear trophoblast markers, KRT7 and CDH1. (C) Box plots of qRT-PCR expression results; bulk rhesus placental samples (purple), iRP cell lines (green). Statistical significance was determined using unpaired t-test (*p<0.05, **p<0.01).

### Transcriptomic comparison of immortalized and primary rhesus trophoblast cells

To characterize gene expression levels in the immortalized trophoblast cell lines and compare them to rhesus primary trophoblast (RPT) cells, RNA-seq was carried out on two replicates each of iRP-D26, iRP-D28A, and freshly isolated first-trimester RPTs. For reference, publicly available human and rhesus peripheral blood mononuclear cells (PBMC) (da Silva Francisco Junior et al. 2019; Micci et al. 2015), human primary trophoblasts (HPT) (Azar et al. 2018), and rhesus bulk placenta (Dunn-Fletcher et al. 2018) RNA-seq datasets were also included in the assessment. Principle component analysis (PCA) based on the expression of all analyzed genes (N=15,787) revealed that a majority of the sample variance was due to tissue-type differences. Both immortalized trophoblast cell line samples (iRP-D26 and iRP-D28A) clustered closely with the RPT samples, highlighting the overall transcriptomic similarity of our newly generated rhesus trophoblast cell lines to freshly isolated RPT cells (Fig. 5A). Moreover, hierarchical clustering of the samples based on the expression of previously defined placental markers showed clustering of iRP-D26 and iRP-D28A with RPT and bulk rhesus placenta samples, as well as HPTs with bulk human placenta, but not with PBMCs from either species (Fig. 5B).

**Figure 5.**
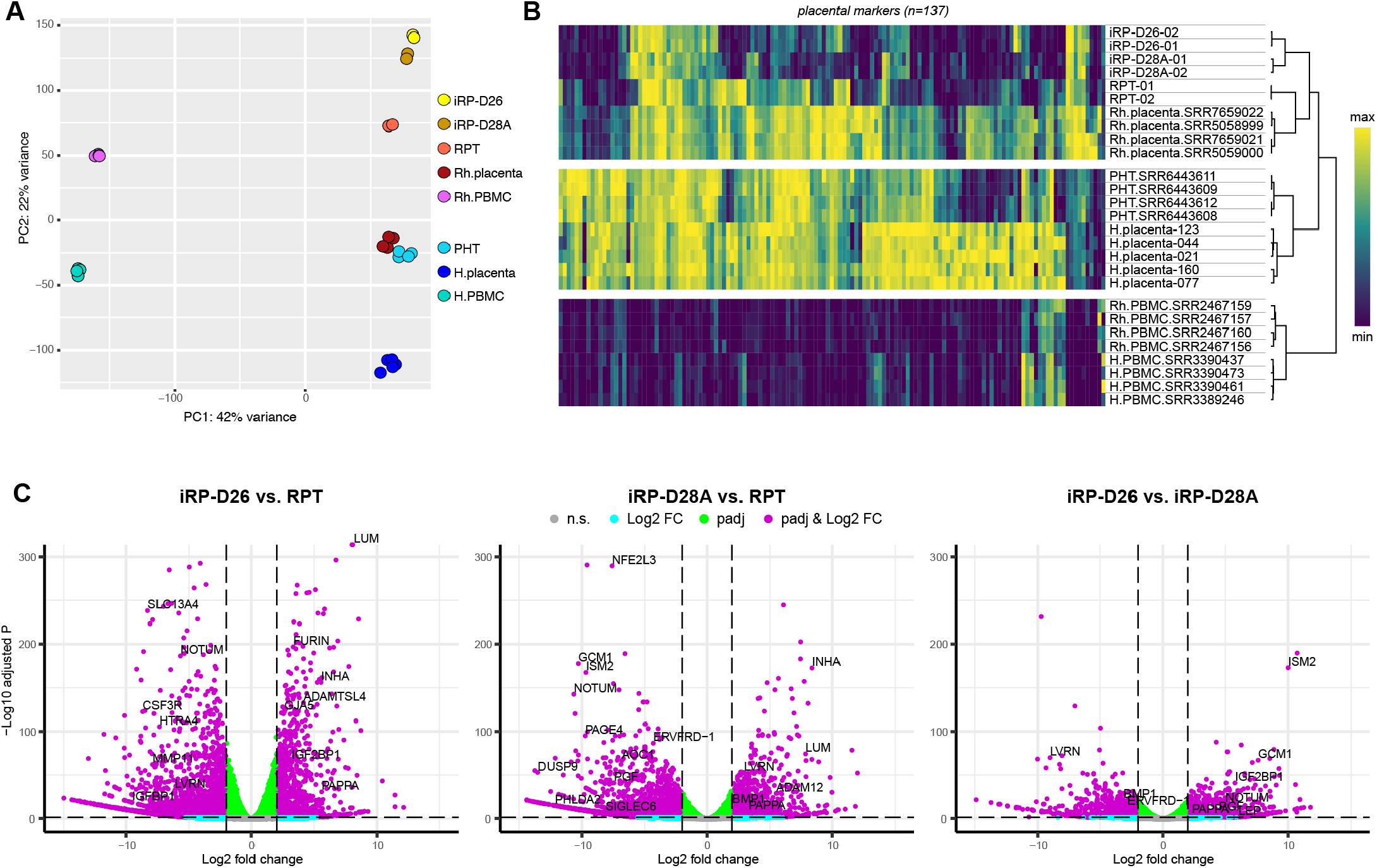
Transcriptomic comparison of immortalized and primary rhesus trophoblast cells. (A) PCA plot of RNA-seq gene expression from human and rhesus bulk placentas, PBMCs, primary trophoblasts, iRP-D26 and iRP-D28A cell lines. (B) Heatmap of placental marker gene expression results from RNA-seq. Color scale depicts minimum (purple) and maximum (yellow) normalized expression values compared across all human and rhesus samples. (C) Volcano plots of DE analysis results; positive Log2FC values represent genes upregulated in iRP-D26 compared to RPT-D50 (left), iRP-D28A compared to RPT-D50 (middle), and iRP-D26 compared to iRP-D28A (right).

Next, we analyzed gene expression differences between the immortalized trophoblast cell lines and RPTs. For this, each immortalized cell line was directly compared to RPTs via two separate DE analyses: (1) iRP-D26 vs. RPT and (2) iRP-D28A vs. RPT. A total of 2927 DEGs (padj<0.05 & Log2FC>|2|) were found between iRP-D26 and RPT, with approximately 70% (N=2055/2927) upregulated in RPT. Further, a total of 2917 DEGs were identified between iRP-D28A and RPT, with approximately 63% (N=1829/2917) upregulated in RPT. Compared to RPTs, both iRP-D26 and iRP-D28A exhibited downregulation of placental genes, *MBNL3* and *NFE2L3* (Fagerberg et al. 2014); and genes involved in immune receptor activity, *LEPR*, *CD74*, *C3AR1*, *CXCR2*, and *CCR* (Raudvere et al. 2019) (Fig. 5C, Supplemental Fig. S4). Upregulation of these genes in primary compared to immortalized rhesus trophoblast cells may be due to the presence of immune cell and/or cytokine-stimulated cells within the freshly isolated RPT samples analyzed. Both iRP-D26 and iRP-D28A showed significant upregulation of lumican (*LUM*), an extracellular matrix proteoglycan highly expressed in trophoblast cells derived from human embryonic stem cells (Yabe et al. 2016), inhibin-α (*INHA*), a member of the transforming growth factor β (TGFβ) superfamily transformative in the differentiation of human CTB to STBs (Azar et al. 2018), and genes involved in cell adhesion (*LAMB3*, *PDCH7*, *SEMA5A*, *CPE*, *LAMA3*, *PTPRD*, and *CDH1*) (Fig. 5C, Supplemental Fig. S4). Further, the iRP-D26 line alone exhibited upregulation of endopeptidase genes, including *MMP1*, *LGMN*, and *FURIN*, a gene required for trophoblast syncytialization (Zhou et al. 2013), whereas iRP-28A showed upregulation of the metalloaminopeptidase genes, *ENPEP* and *LVRN*, markers of trophoblast progenitors (Krendl et al. 2017) and EVTs (Fujiwara et al. 2004), respectively (Fig. 5C, Supplemental Fig. S4). These findings suggest that the iRP-D26 line was comprised of villous CTB-like progenitor cells that normally fuse to form STBs, whereas the iRP-D28A line was more EVT-like in origin.

### iRP-D26 and iRP-D28A express genes consistent with undifferentiated CTB progenitor cells

While there are a number of established markers for human CTBs, EVTs, and STBs, most have not been validated in rhesus and thus, an equivalent set of high-confidence rhesus trophoblast markers is currently lacking. To identify rhesus trophoblast subtype markers and determine if they were expressed in our cell lines, we first assessed the expression levels of several genes throughout *in vitro* differentiation of RPT cells. Freshly isolated RPT cells were allowed to spontaneously differentiate and fuse in culture for 72 hours. Samples were collected at 24hr intervals throughout differentiation and analyzed for fusion via IF staining and gene expression via qRT-PCR. IF staining of the membrane marker, CDH1, was used to monitor differentiation and fusion of RPT cultures over time. At 24hrs, RPT cultures contained abundant CDH1 positive mononuclear cells and some small regions of multinucleated STB. By 48hrs and 72hrs, CDH1 staining was drastically diminished and the RPT cultures predominantly consisted of large multinucleated STB cells (Fig. 6A). Following the natural progression of trophoblast fusion, these results suggest that proliferative and fusogenic CTBs are most highly represented at the earliest time points (0hrs and 24hrs), while differentiated multinucleated STBs primarily appear at the latest time points (48hrs and 72hrs).

**Figure 6.**
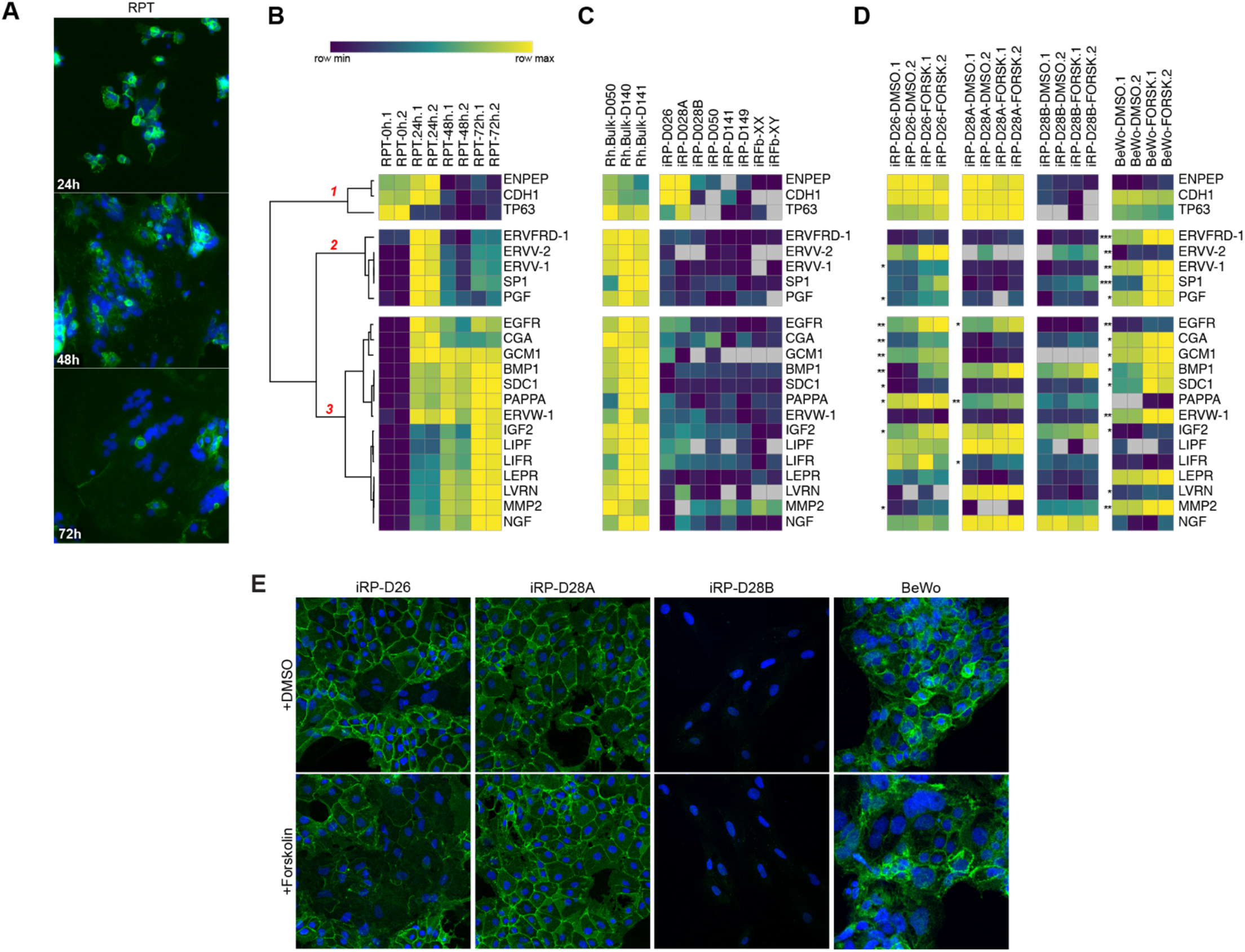
Analysis of cell fusion and STB differentiation in primary and immortalized rhesus trophoblast cells. (A) CDH1 IF staining of PRT cells after 24hrs, 48hrs, and 72hrs in culture. (B-D) Heatmaps of qRT-PCR RNA expression levels. Color scale depicts minimum (purple) and maximum (yellow) Log2 relative gene expression values compared across (B) RPT cells after 24hrs, 48hrs, and 72hrs in culture; the three gene clusters identified via hierarchical clustering are labeled in red. (C) bulk rhesus placenta, iRP, and iRFb cell lines; (D) iRP and BeWo cell lines treated with DMSO or Forskolin for 48hrs. Asterisks denote significant expression difference between DMSO and forskolin treated samples; *P-value<0.05, **P-value <0.01, ***P-value <0.001. (E) IF staining of iRP and BeWo cell lines treated with DMSO or Forskolin for 48hrs; multinucleated cells are observed in both iRP-D26 and BeWo cell lines upon forskolin treatment; CDH1 (green) and DAPI (blue).

The IF findings were further supported by hierarchal clustering of qRT-PCR expression results, which revealed three clusters of genes based on RPT differentiation time points (Fig. 6B). Cluster 1 contained genes with high expression in freshly isolated RPT samples (0hrs) and included well-known human trophoblast progenitor markers such as *ENPEP* (Krendl et al. 2017) and *TP63* (Renaud et al. 2015; Soncin et al. 2018). Cluster 2 expression levels were high only at 24hrs, and included several genes associated with fusogenic CTBs in primates: *ERVFRD-1* (Esnault et al. 2008), *ERVV-2*, and *ERVV-1* (Esnault et al. 2013; Kjeldbjerg et al. 2008). Cluster 3 genes showed sustained or increased expression levels in the later time points (48hrs and 72hrs), and included several established markers of human differentiated STB cells: *GCM1* (Liang et al. 2010), *BMP1* (Aronow et al. 2001), *SDC1* (Crescimanno et al. 1999), *PAPPA* (Guibourdenche et al. 2003), and *ERVW-1* (Blond et al. 2000). A few genes associated with human EVT cells were also noted within cluster 3, including *LIFR* (Tilburgs et al. 2015), *LVRN* (Horie et al. 2012), and *MMP2* (Onogi et al. 2011). A similar analysis of the immortalized rhesus cell lines revealed a high expression level of cluster 1 genes (*ENPEP*, *CDH1*, and *TP63*) in iRP-D26 and iRP-D28A cells. Thus, both the iRP-26 and iRP-D28A cell lines appeared more similar to undifferentiated proliferative CTB progenitor cells than their differentiated trophoblast counterparts (Fig. 6C).

### Forskolin induces STB differentiation in iRP-D26 trophoblast cells

To test the fusogenic potential of iRP-D26 and iRP-D28A we treated the cell lines with forskolin, an activator of adenylate cyclase and known inducer of fusion and STB formation in BeWo human choriocarcinoma (Wice et al. 1990) and trophoblast stem cells (Okae et al. 2018). Human BeWo and iRP-D28B (largely comprised of mesenchymal cells) were included as positive and negative controls, respectively. Each of the cell lines was treated with 25uM of forskolin or DMSO (vehicle control) for 48hrs, then analyzed for gene expression changes in cluster 1-3 genes via qRT-PCR. Compared to DMSO, forskolin treated BeWo cells showed a significant increase in fourteen of the genes tested. These included all cluster 2 genes and many of cluster 3 genes, primarily corresponding to fusogenic CTB and STB genes, respectively (Fig. 6D). Similar results were obtained for iRP-D26 forskolin treated cells, which showed a significant upregulation of all genes identified in BeWo, except *ERVFRD-1*, *ERVV-2*, *SP1*, *ERVW-1*, and *LIFR* (Fig. 6D). Forskolin treatment of iRP-D28A cells showed a significant increase in only three of the genes tested (*EGFR*, *PAPPA*, and *LIFR*), while no significant expression changes were identified in forskolin treated iRP-D28B cells used as a negative control (Fig. 6D). Consistent with qRT-PCR expression results, IF analysis showed no changes in multinucleation or CDH1 staining between DMSO and forskolin treated iRP-D28A or iRP-D28B cells. However, IF staining of both iRP-D26 and BeWo forskolin treated cells revealed a notable increase in multinucleated cells and a corresponding reduction in CDH1 staining (Fig. 6E). Collectively, these results highlighted the fusogenic potential of the iRP-D26 cell line and further suggested that the iRP-D28A cells were more EVT-like.

## Discussion

The wide variety of placental morphologies and physiologies that exist among mammals makes it difficult to adequately model human placentation and placental pathologies in other species (Mossman 2013; Cross et al. 2003; Carter and Enders 2004). However, many distinctive features of human placentation are reportedly conserved in rhesus monkeys (Ramsey and Harris 1966; Ramsey et al. 1976; Ellinwood et al. 1989; King and Blankenship 1994; Blankenship and Enders 2003). Despite this conservation, no study has been conducted to comprehensively assess the molecular similarities and potential differences between human and rhesus placental tissues. Comparative analyses of human and closely-related species are beginning to identify specific genetic and molecular changes that seem to account, in part, for specific aspects of human evolution, including human diseases (Rogers and Gibbs 2014). Thus, identification of the molecular differences between human and rhesus placenta is not only needed to elucidate the translatability between human and rhesus placental studies, but it may also provide valuable insight into the molecular origin of human placental diseases, such as pre-eclampsia. Here, we performed a cross-species transcriptomic comparison of human and rhesus placental tissue in order to identify molecular differences and ultimately elucidate the translatability between human and rhesus placental investigations. Further, to increase the accessibility of rhesus *in vitro* placental studies, we generated and thoroughly characterized two highly-pure TERT-immortalized rhesus trophoblast cell lines that retained features of primary rhesus trophoblasts cells.

Differential expression analysis of human and rhesus placental tissue revealed that while a majority of human placental marker genes were similarly expressed, certain genes were differentially expressed between the two species. Specifically, several genes associated with invasive EVTs and pregnancy complications were shown to be upregulated in human compared to rhesus placenta. These results are consistent with previously reported differences between human and rhesus placentation, such as the increased level of EVT invasion (Ramsey and Harris 1966; Ramsey et al. 1976; Pijnenborg et al. 2011) and heightened risk of pre-eclampsia in humans (Rosenberg and Trevathan 2007). However, the majority of the DEGs identified here have not been previously reported, thus our results provide novel insight into the molecular differences underlying human and rhesus placentation. For instance, we observed the upregulation of *LGALS13* in human compared to rhesus, which encodes a galectin that interacts with glycoproteins and glycolipids to facilitate the expansion of uterine arteries and veins during pregnancy in an endothelial cell-dependent manner via the eNOS and prostaglandin signaling pathways (Sammar et al. 2019). Its downregulation in pregnancy disorders is thought to contribute to aberrant uteroplacental blood flow (Sammar et al. 2019), while the decreased expression in rhesus placenta may underlie differences in the extent and depth of human versus rhesus placental invasion or perhaps represent an alternative mechanism for uterine vessel expansion in rhesus. Our results also suggest that recently-evolved highly-expressed human placental genes may contribute to an increased risk of aberrant EVT invasion associated with pre-eclampsia (Brosens et al. 2011); however, further evolutionary and functional investigation of these genes are required to determine this.

Approximately 14% (N=61/447) of the genes we identified as upregulated in human were previously annotated in other comparative studies as specifically expressed or upregulated in human placenta compared to many distantly-related species (Hou et al. 2012; Armstrong et al. 2017). However, since these studies relied on the comparison of placental gene expression across numerous distantly-related species, they were limited in the number of orthologous genes eligible for differential expression comparison. Overall, only ~4% of the genes we identified as upregulated in human were also identified by Armstrong et. al (*ADAM12, GULP1, SERPINB2, CYP19A1, MUC15, SVEP1, GPC3, FBN2, DEPDC1B, ATG9B, EFHD1, KISS1, FAM13A, PKIB, S100A9, SCIN, OLAH, CCK*), suggesting that the expression of these genes is highly divergent not only between human and rhesus but also between human and other species. Two of these genes, *ADAM12* and *KISS1*, are associated with pre-eclampsia (Enquobahrie et al. 2012; Kapustin et al. 2020) supporting our finding that pre-eclampsia related genes are significantly over-represented in the human-upregulated placental gene set.

Since the placenta contains many cell types in addition to trophoblasts (Liu et al. 2018), primary trophoblast cells were isolated from bulk rhesus placental for further characterization, as well as TERT-immortalization, to generate a previously unavailable resource for functional testing of first-trimester rhesus trophoblasts cells. In total, six TERT-immortalized rhesus trophoblast cell lines were generated, however, only two of the lines (iRP-D26 and iRP-D28A) were highly pure and devoid of large-scale CNVs despite small sub-chromosomal changes. These results are consistent with previous studies that revealed few karyotypic differences in TERT-immortalized cells compared to SV40-immortalized cells (Toouli et al. 2002). Nevertheless, genome duplication may still occur in TERT-immortalized trophoblast cells over time with continued passaging (Reiter et al. 2017), suggesting that the genome integrity of TERT-immortalized cell lines should be routinely monitored. We note that previously generated human trophoblasts have been shown to preserve the phenotypic properties that closely resemble *in vivo* characteristics (Wieser et al. 2008; Straszewski-Chavez et al. 2009). Through a combination of DE, qRT-PCR, and forskolin-stimulation analyses, we found iRP-D26 specifically retained features of villous CTBs, while iRP-D28A retained features of cell column CTBs, progenitor cell populations known to differentiate into STBs and EVTs, respectively (James et al. 2005; Gamage et al. 2016).

When grown in culture, primary human trophoblast cells spontaneously differentiate and fuse into STBs (or a syncytial layer) over time (Kliman et al. 1986). Similar to what is observed with human, we found that after 72hrs in culture, almost all RPT cells fused into multinucleated syncytia. Additionally, known human STB markers (*GCM1* and *PAPPA*) were found to increase with RPT differentiation, whereas expression of the human fusogenic CTBs marker (*ERVFRD-1*) peaked at 24hr and decreased with RPT differentiation. Upon comparison to the immortalized trophoblasts, both cell lines retained high expression levels of known mononuclear trophoblast cell markers (Lee et al. 2016). However, compared to RPT cells, iRP-D26 showed upregulation of *FURIN*, a gene required for trophoblast syncytialization (Zhou et al. 2013). In contrast, iRP-28A showed upregulation of the metalloaminopeptidase genes, *ENPEP* and *LVRN*, markers of trophoblast progenitors (Krendl et al. 2017) and EVTs (Fujiwara et al. 2004), respectively. While there are previous reports of rhesus trophoblast stem cell lines derived from blastocyst outgrowths (VandeVoort et al. 2007; Matsumoto et al. 2020), we are the first to describe the generation and characterization of sustainable rhesus first-trimester trophoblast cell lines isolated from placental tissue. Our data suggest that these newly generated TERT-immortalized rhesus trophoblast cell lines represent a useful *in vitro* model for future primate trophoblast studies.

In conclusion, our comparative analysis between human and rhesus bulk placenta showed that while a majority of placental marker genes are similarly expressed between the two species, certain genes are differentially expressed between human and rhesus placenta. Moreover, we generated immortalized rhesus trophoblast cell lines that represent a useful tool for future primate placental investigations, especially for *in vitro* experiments that interrogate the putative function of genes identified in this study. Transcriptomic comparison and functional assessment of these cell lines suggest that they retain attributes of primary first-trimester rhesus trophoblasts required for additional trophoblast cell differentiation studies. Collectively, these results suggest that rhesus is a suitable surrogate for most investigations of human placentation, however, notable molecular differences related to EVT function, pre-eclampsia and/or other pregnancy disorders should be considered and further interrogated in future investigations.

## Methods

### Tissues and cell lines

Deidentified human term placental samples were collected by and acquired through the Labor and Delivery Unit at the Oregon Health and Science University Hospital and deposited into a repository under a protocol approved by the Institutional Review Board with informed consent from the patients. A total of five different human placentas from healthy cesarean section term births, ranging from 38.9 to 41.3 gestational weeks, were used for RNA-seq library generation (Supplemental Table S1). All rhesus monkey (*Macaque mulatta*) tissues were collected in compliance with the guidelines established by the Animal Welfare Act for housing and care of laboratory animals and conducted per the Institutional Animal Care and Use Committee (IACUC protocols #0514 and #0580) at the Oregon National Primate Research Center (ONPRC). All rhesus placentas were collected from time mated breeding pregnancies delivered via cesarean section. Two frozen rhesus third-trimester placental samples, collected at 140 and 141 gestational days, were used for RNA isolation and qRT-PCR validation of DE analysis results. Six fresh rhesus placentas were used for primary rhesus trophoblast TERT-immortalization, including two term placentas (D141, D149) and four first-trimester placentas (D26, D28A, D28B, D50). An additional first-trimester rhesus placenta (D50) was used for primary trophoblast culture and RNA-seq analysis; these cells were not included in TERT-immortalization experiments. For these samples, the placentas were separated from the fetus and amniotic sac, collected in cold sterile saline and immediately processed for isolation of primary trophoblasts. The primary female and male rhesus macaque skin fibroblasts cell lines, Fb.XX (AG08312) and Fb.XY (AG08305), were acquired through Coriell Institute.

### RNA isolation and purification

Frozen placental samples were ground into a powder using liquid nitrogen-cooled mortar and pestle then directly added to TRIzol reagent (Thermo Fisher #15596026); for cell lines media was removed and TRIzol reagent was added directly to the tissue culture dish. RNA was isolated from TRIzol reagent, treated with Turbo DNAse (Thermo Fisher #AM1907), and purified using RNA Clean and Concentrator-5 spin columns (Zymo #R1013) according to manufactures instructions.

### RNA-seq library preparation and data acquisition

NEBNext® Ultra II Directional RNA Library Prep Kit for Illumina and NEBNext rRNA Depletion Kit (NEB, Ipswich, MA) was used to generate RNA-seq libraries from purified RNA following the manufacturer’s instructions. Libraries were quantified with the Qubit High Sensitivity dsDNA Assay (Invitrogen, Carlsbad, CA) and size distribution was assessed with a 2100 Bioanalyzer High Sensitivity DNA Analysis Kit (Agilent). Multiplexed bulk human placental libraries were sequenced on the NextSeq500 platform using 150 cycle single-end protocol generating a total of 36.9 to 70.9 million 101bp reads per sample. Multiplexed rhesus trophoblast cell lines were sequenced on the NextSeq500 platform using 100 cycle single-end protocol generating a total of 57.8 to 68.5 million 75bp reads per sample. Additional publicly available term placental RNA-seq data from human and rhesus were downloaded from NCBI SRA using SRA Toolkit (http://ncbi.github.io/sra-tools/); rhesus placenta: SRR5058999, SRR5059000, SRR7659021, SRR7659022; human placenta (DE#2): SRR3096525, SRR3096545, SRR3096594, SRR3096612, SRR3096624, SRR3096625; human placenta (DE#3): SRR4370049, SRR4370050; human PBMC: SRR3389246, SRR3390437, SRR3390461, SRR3390473; rhesus PBMC: SRR2467156, SRR2467157, SRR2467159, SRR2467160; HPT: SRR6443608, SRR6443609, SRR6443611, SRR6443612 (Supplemental Table S1).

### Human and rhesus orthologous gene annotations

Human protein-coding gene annotations (GRCh38.99) including associated rhesus orthologous gene annotations (Mmul10.99) were downloaded from Ensembl BioMart (Zerbino et al. 2017; Kinsella et al. 2011). Gene annotations used for differential expression analysis were filtered to include only protein-coding genes with high-confidence “one2one” rhesus orthologous genes, producing a final set of 15,787 human gene annotations and associated 15,787 rhesus orthologs (Supplemental Table S6). A total of 13,478 orthologues genes passed the minimum DEseq2 default expression threshold for differential expression statistical analysis.

### Differential expression analysis

Raw fastq files were trimmed of low-quality and adapter sequences using Trimmomatic (Bolger et al. 2014) and mapped to both the human (GCh38) and rhesus (Mmul10) reference genomes using Bowtie2(Langmead et al. 2019) with --very-sensitive parameter. Resulting BAM files were filtered to remove low quality and multi-mapped reads (MAPQ<10) using samtools (Li et al. 2009) view -q 10. Raw read counts for GRCh38.99 human gene annotations were generated from GRCh38 mapped data, while raw read counts for Mmul10.99 rhesus gene annotation were generated from Mmul10 mapped data, using featureCounts (Liao et al. 2014)–primary and filtered to include gene annotations described above. Gene counts were normalized and differentially expressed genes (DEGs) (padj<0.05 & Log2FC>|2|) were identified using default setting of DEseq2 (Love et al. 2014). DE analysis was performed with human mapped data (DE-GRCh38) and with rhesus mapped data (DE-Mmul10). A gene was considered differentially expressed only if it was identified as significantly (padj<0.05) upregulated or downregulated (|L2FC|>2) by both DE-GRCh38 and DE-Mmul10 analyses. The DE analysis was repeated a total of three times, with three independent sets of human placental RNA-seq data (Supplemental Fig. S1A-C). The first DE analysis included the five human placental RNA-seq samples generated by our group (DE#1), the second DE analysis included six publicly available human placental RNA-seq datasets (DE#2), and the third DE analysis included two publicly available human placental RNA-seq datasets (DE#3); all three DE analyses included the same four publicly available rhesus placental RNA-seq datasets. The final set of DEGs consisted only of genes determined to be significantly upregulated or downregulated by all three DE analyses. The DE analysis of iRP and RPT cells was performed as described above for rhesus (Mmul10) mapped data. A total of 15,787 genes were analyzed, and genes were identified as significantly differentially expressed if the adjusted p-value was less than 0.05 (padj<0.05). For PCA and heatmap visualizations, all datasets were mapped to the rhesus genome (Mmul10), and DEseq2 variance stabilizing transformation was used to normalize raw gene counts. All 15,787 analyzed genes were included in PCA. Morpheus webtool (https://software.broadinstitute.org/morpheus) was used to generated heatmap and perform hierarchical clustering (metric: one minus pearson correlation, linkage method: complete). Statistical over-representation analysis of the human and rhesus upregulated gene lists was performed using g:Profiler (Raudvere et al. 2019) with a background gene list containing the final gene annotations list described above. In addition to the default Gene Ontology (GO) gene sets supplied by the g:Profiler webtool, “Jensen_DISEASES” and “Rare_Diseases_AutoRIF_Gene_Lists” human disease annotation gene set libraries were downloaded from the Enrichr webtool and analyzed using g:Profiler to identify over-representation of disease-associated genes. Functional gene sets were identified as significantly over-representation if pad<0.05.

### Quantitative reverse transcriptase PCR (qRT-PCR)

Primers were carefully designed to amplify both human and rhesus sequences of all genes examined, with noted exceptions (Supplemental Table S7). Purified RNA was reverse transcribed into complementary DNA (cDNA) using SuperScript VILO cDNA Synthesis Kit (Thermo Fisher #11754050). Samples were prepared for high-throughput qRT-PCR using 96.96 gene expression dynamic array (Fluidigm BioMark) following manufactures protocol “Fast gene expression analysis using Evagreen”. Briefly, preamplification of cDNA was performed using 500nM pooled primer mix, unincorporated primers were removed with exonuclease I treatment, and diluted 5-fold before samples and detectors were loaded and run on a 96.96 array with the following thermocycler settings: 70°C for 40 min., 60°C for 30 sec., 95°C for 1 min., 40 cycles of 95°C 15 sec., 59.5°C for 15 sec, 72°C for 15 sec. Two no template control (NTC) samples and four technical replicates of each sample-detector combination and were included. The analysis was performed using qbase+ software, with GAPDH, HPRT1 and TBP serving as the reference genes used for normalization. Statistical significance was determined for each gene/detector using unpaired t-tests. Mean calibrated normalized relative quantities (CNRQ) were exported from qbase+, Log2 transformed, then used as input for heatmap generation with Morpheus web tool.

### Human placental marker gene set analysis

Human placental markers were defined by combining previously identified placenta “tissue enriched” and “group enriched” genes from The Human Protein Atlas (Uhlén et al. 2015). Placenta enriched genes were identified by having at least four-fold higher mRNA level in the placenta compared to any other tissue, while placenta group enriched genes were identified by having at least four-fold higher average mRNA level in a group of 2-5 tissues compared to any other tissue (Uhlén et al. 2015). Ensemble GRCh38.99 rhesus homology details were used to identify human placental marker genes without a known rhesus ortholog, lacking a high-confidence rhesus ortholog, or not defined as a one2one ortholog; since our DE analysis relies on the comparison of human and rhesus orthologous genes, these markers could not be included. The results from our DE analysis were used to identify placental markers with altered expression levels between human and rhesus, and classified as either human-upregulated, rhesus-upregulated, or not differentially expressed.

### Primary rhesus trophoblast cell isolations

Primary trophoblast cells were isolated from rhesus placentas using protocols adapted from previously described methods for human first-trimester tissue (Straszewski-Chavez et al. 2009) and human term tissue (Kliman et al. 1986). All rhesus placentas were obtained immediately after cesarean section delivery, and procedures were performed in a biosafety cabinet using ice-cold and sterile solutions unless otherwise noted. Placental tissue was transferred to a petri dish and covered with sterile saline, and the villous tissue was dissected from the decidua and chorionic plate using scissors and forceps; decidua and fetal membranes were discarded. To remove any contaminating blood, the villous tissue was washed until clear with several changes of sterile saline then crudely minced using scissors. For first-trimester placentas, villous tissue was transferred to a 50ml tube containing warmed 0.25% trypsin solution and incubated at 37°C for 10 min, inverting mixing every 2-3 min. The supernatant containing STBs and EVTs was discarded and the tissue was washed with three changes of 1X phosphate-buffered saline without Ca^2+^ and Mg^2+^ (PBS^−^) (Caisson Labs #PBL05). To release the CTB, the tissue was transferred to a fresh petri dish containing warmed 0.25% trypsin 0.2mg/mL DNAse I solution and a scalpel or glass slide was used to thoroughly scrape the villi. The surrounding trypsin solution containing desired CTBs was collected through a 70μm cell strainers into 50ml tubes containing 5ml fetal bovine serum (FBS) (Fisher #16-140-063). Cells were centrifuged at 300g for 10min and supernatants were disposed of. The cell pellet was resuspended cell culture media (CCM): DMEM high-glucose glutaMAX (Fisher, #10566-016), 10% FBS, 100 U/ml Pen-Strep (Fisher #15-140-148); carefully layered on top of 5ml lymphocyte separation media (LSM) (Corning #25-072-CI) in a 15ml tube, and centrifuged at 400g for 15min with break off. The mononuclear layer at the LSM-media interface was carefully collected into a new 15ml tube, counted, and washed with maximum volume of CCM. For term placentas, villous tissue was digested for 30 minutes shaking at 37°C with 0.25% trypsin and 0.2mg/mL DNAse I. Supernatant was reserved and the digest was repeated two additional times. The three digests were combined and centrifuged through Normal Calf Serum. Pellets were resuspended in DMEM and re-pelleted. Cells were layered over a Percoll gradient and spun at 2800rpm for 30 minutes without brake. The CTB cells between 35% and 55% were collected, counted and resuspended in CCM. For both first-trimester and term placentas, cells were centrifuged at 300g for 10 minutes, and the cell pellet was resuspended to 10^8^ cells/mL in 1X Nanobead buffer (BioLegend #480017). Contaminating immune cells were depleted using anti-CD45 Magnetic Nanobeads (BioLegend #488028) following the manufactures instructions. Purified trophoblast cells were resuspended in complete trophoblast media (CTM): MEM – Earle’s with D-Val (Caisson Labs #MEL12), 10% Normal Human Serum (Gemini Bio #100-110), 100 U/ml Pen-Strep, 1mM Sodium Pyruvate (Fisher #11-360-070), and 0.1M HEPES (Fisher #15-630-106); and grown on enhanced tissue culture dishes (Corning Primaria, #C353802) in a humidified 37°C environment with 5% CO_2_. Primary rhesus trophoblasts included in RNA-seq (D50B) were harvested after nanobead immuno-purification and additional CTM wash step.

### TERT-immortalization

Primary rhesus trophoblasts and rhesus skin fibroblast cells were immortalized using Alstem’s TERT-immortalization kit (Alstem #CILV02) following manufactures instructions. In brief, following isolation of primary cells or 24hr after thawing of skin fibroblasts, the cells were plated at a density of 1.5 × 10^5^ cells/well in 6-well plate and transduced the following day. Each well received 1mL of media containing 4uL recombinant TERT lentivirus and 500x TransPlus reagent (Alstem # V020). After 16 hours, the media was replaced with fresh culture media and the cells were allowed to recover for 48 hours before beginning puromycin selection (Santa Cruz Biotechnology #SC-108071). Cells were treated with 800ng/ml puromycin for a total of 72 hours. The surviving cells were propagated and represent the TERT-immortalized cell lines established and characterized throughout these studies. Mock transductions using the same transduction conditions without lentivirus added to media were included throughout puromycin selection to ensure depletion of non-transduced cells.

### Cell culture and forskolin treatment

All cell lines were grown in a humidified 37°C environment with 5% CO_2_. Cell culture media was changed every 2 days, and cells were enzymatically passaged using TrypLE (Gibco). Primary and TERT-immortalized trophoblast cell lines were cultured in CTM, while fibroblast samples were cultured in CCM. D26 and D28A cell lines were passaged at a density of 30,000 cells/cm^2^, while all other lines were passaged at a density of 15,000 cells/cm^2^. For forskolin treatment, the cells were passaged onto 6-well plates and treated the following day with either media containing 25uM forskolin or DMSO (vehicle-control); the cells were harvested 48 hours later for analysis.

### DNA sequencing and chromosome copy number calling

Cells were dissociated using TryplE, pelleted, and resuspended in PBS^−^ containing 0.05% trypsin-EDTA (Thermo Fisher Scientific). A stereomicroscope was used to isolate, wash, and collect cells into individual sterile PCR tubes. Immediately after collection, PCR tubes containing single-cells (n=4), five-cells (n=1), and ten-cells (n=1) were flash-frozen on dry ice, and stored at −80C until library preparation. Individual samples underwent DNA extraction and whole-genome amplification, library pooling, DNA sequencing were performed as previously described (Daughtry et al. 2019). Multiplexed libraries were loaded at 1.6pM and sequenced on the NextSeq 500 platform using 75 cycle single-end protocol. The resulting sequencing data was filtered, trimmed and mapped to the rhesus reference genome (Mmul8) as previously described (Daughtry et al. 2019). Chromosome copy number calling and plots were generated using Ginkgo (Garvin et al. 2015). The proportion of ChrY reads was determined for each sample using by dividing the number of reads mapped to ChrY by the total number of mapped reads. The relative proportion of ChrY reads was identified by normalizing samples to the known male sample (iRFb-XY), and samples with a mean relative proportion ≥ 0.50 were identified as male. Box and whisker plots depicting mean (dashed line) and median (solid line) and were generated using Plotly chart studio.

### Metaphase spread chromosome counts

Cells were treated with 0.015ug/ml colcemid overnight (~12 hours) to induce metaphase arrest. Cells were dissociated using TryplE, pelleted, and resuspended in a warm hypotonic solution (0.06 M KCl, 5% FBS) for 15 minutes before being fixed with 3:1 methanol:acetic acid. Slides were made and baked at 95C for 20 minutes, cooled, trypsinized for 45 seconds and stained with Wright’s stain. At least 20 metaphase spreads brightfield images were captured using 100X objective on Nikon microscope and counted using FIJI software. Box and whisker plots depicting mean (dashed line) and median (solid line) and were generated using Plotly chart studio.

### Immunofluorescence (IF)

The cell culture media was removed from cells, fixed with ice-cold methanol for 15 minutes at −20C, then washed in three changes of PBS. Nonspecific proteins were blocked by incubating cells in 5% donkey serum for 30 min. anti-KRT7 (mouse monoclonal, Dako, M7018), anti-CDH1 rabbit monoclonal (Cell Signaling, 3195S), anti-VIM rabbit monoclonal (Cell Signaling, 5741T), and anti-PTPRC rabbit monoclonal (Cell Signaling, 13917S) antibodies were diluted in PBS + 1% BSA (KRT7 1:250, CDH1 1:250, VIM 1:250, CD45 1:250), and incubated for 2hrs at room temperature. The cells were then washed in three changes of PBS, before incubating in secondary antibody dilutions for 1 hour at room temperature. Alexa Fluor 488 and 594 (Life Technologies) secondary antibody dilutions were prepared at 1:1000 in PBS + 1% BSA. Cells were counterstained with DAPI and washed with three changes of PBS before imaging. Images of cells were captured using 20X objective on epifluorescence microscope and processed using FIJI software.

### Immunohistochemistry (IHC)

Paraffin sections were deparaffinized and rehydrated through xylene and graded alcohol series, then washed for 5 min in running tap water. Antigen unmasking was performed using sodium citrate (pH 6.0) buffer in a pressure cooker for 20 min, washed in three changes of PBS. An endogenous enzyme block was performed by incubating sections in 0.3% hydrogen peroxide for 10 min, washed in three changes of PBS. Nonspecific proteins were blocked by incubating sections in 5% horse serum for 30 min. Primary antibodies were diluted as described for IF staining, and tissue was incubated in primary antibody dilutions for 2hrs at room temperature. Mouse IgG H+L (Vector Labs, BA-2000) and rabbit IgG H+L (Vector Labs, BA-1100) biotinylated secondary antibody dilutions were prepared at 1:250 in PBS + 1% BSA. The tissue sections were incubated in secondary antibody dilution for 1hr at room temperature, then washed in three changes of PBS. VECTASTAIN Elite ABC HRP Kit (Vector Laboratories, PK-6100) and ImmPACT DAB Peroxidase HRP substrate (Vector Labs, SK-4105) were used according to manufactures instructions. Nuclei were counterstained with hematoxylin (Electron Microscopy Sciences, 26043-05), and imaged using a brightfield microscope.

## Data access

All RNA-seq and DNA-seq raw sequencing data generated in this study have been submitted to the NCBI Sequencing Read Archive (SRA; https://www.ncbi.nlm.nih.gov/sra/) under BioProject accession number PRJNA649979.

## Acknowledgements

We thank L. Myatt for access to human placental samples; J. Hennebold and A. Clark for access to rhesus first-trimester placental samples; A. Frias for use of tissue culture facilities and access to rhesus third-trimester primary trophoblast cells; A. Adey and A. Fields for the assistance and use of NextSeq500 sequencer; J. Hennebold and M. Murphy for use of tissue culture and microscope facilities; S. Stadler, A. Adey, J. Sacha, M. Okhovat, and B. Davis for insight and advice; and The Molecular Technologies Core at ONPRC. J.L.R was funded by the NIH/NICHD grant no. F31HD094472.

## Author contributions

J.L.R, S.L.C. and L.C. designed research; J.L.R, J.G. and V.R performed research; J.L.R. analyzed data; S.L.C. and L.C. supervised progress of all research; and J.L.R, S.L.C. and L.C. wrote the paper. All authors were involved in manuscript editing.

